# PP2A forestalls mitophagy by dephosphorylating Parkin and ubiquitin

**DOI:** 10.1101/2022.03.12.484070

**Authors:** Shang-Xiang Ye, Xing-Yu Liu, Ze-Feng Nie, Ling-Shen Meng, Xu Dong, Fang-Fang Li, Zhou Gong, Wei Yang, Wei-Ping Zhang, Chun Tang

## Abstract

Mitophagy is a selective autophagic process that removes damaged mitochondria. PINK1-Parkin axis is primarily responsible for initiating mitophagy via feedforward mechanism, in which PINK1 phosphorylates ubiquitin and Parkin, and Parkin gains E3 ligase activity at the mitochondrial outer membrane. However, the phosphatase of pParkin is unknown, and the braking mechanism of mitophagy is incomplete. Here we report that protein phosphatase 2A (PP2A) catalyzes the dephosphorylation of Parkin and ubiquitin, with Cα, Aα, and B55α, the catalytic scaffolding and regulatory subunits, respectively. Up- or down-regulation of PP2A protein level in cells by over-expression or transfection of specific siRNA decreases or increases Parkin and ubiquitin phosphorylation levels, respectively. Consequently, PP2A phosphatase activity negatively modulates mitochondrial translocation of Parkin and forestalls mitophagy. Our finding thus places PP2A, an enzyme already known for its involvement in the pathogenesis of neurodegenerative diseases, further intertwined with the PINK1-Parkin signaling pathway.

---

Mitophagy removes damaged mitochondria through selective autophagy, and preserves the overall quality and quantity of mitochondria. PINK1, a kinase lodged at the outer mitochondrial membrane (OMM), usually undergoes rapid turnover^1,2^, but becomes accumulated and activated at the sites with depolarized mitochondrial membrane potential^3-5^. Once activated, PINK1 phosphorylates ubiquitin at residue S65^6-9^. The phosphorylated ubiquitin (pUb) has a higher affinity for Parkin than the unmodified ubiquitin does^7,10,11^, and thus recruits Parkin, a promiscuous E3 ligase, to OMM, which can eventually lead to mitophagy.

The activation mechanism of Parkin has been well established. Briefly, the binding of pUb to Parkin unhinges Parkin N-terminal Ubl domain allosterically. Once dislodged, Parkin Ubl domain becomes more accessible and can be readily phosphorylated by PINK1, also at residue S65^11-13^. PINK1 phosphorylation stabilizes Parkin Ubl domain in the dislodged flexible state, leads to a further unraveling of the Parkin RING2 domain, and fully enables Parkin E3 ligase activity^10-12,14,15^. Once activated, Parkin ubiquitinylates a multitude of OMM proteins related to mitophagy^16-19^. Thus, through tight regulation of Parkin activation, the PINK1-Parkin axis constitutes the primary pathway for mitophagy initiation.

Conversely, negative regulatory mechanisms of mitophagy have been reported. Deubiquitinases including USP14, USP15, and USP30 are known to remove polyubiquitin modifications from the substrate proteins of Parkin and antagonize Parkin-mediated mitophagy^20-22^. More recently, PTEN-L, a translational variant of PTEN, and PPEF2, abundant at the inner segment of rod cells and known for its role in sensory transduction^23^, have been shown as the phosphatases for pUb^24-26^. The pUb can be involved in a multitude of cellular activities beyond mitophagy^13,27,28^. Considering the central role of Parkin in mitophagy, dephosphorylation of Parkin is a more specific and efficient way to keep mitophagy in check. However, the phosphatase of pParkin has not been unambiguously identified.

With conjoint use of nuclear magnetic resonance (NMR) and biochemical analyses, here we report that protein serine/threonine phosphatase 2A (PP2A) dephosphorylates both pParkin and pUb. A PP2A holoenzyme comprises a catalytic subunit C, a scaffolding subunit A, and a variable regulatory subunit B^29-31^. Cα and Aα are the predominant catalytic and scaffolding isoforms of PP2A, respectively, while the regulatory subunits fall into four families, including B (also known as B55 or PR55), B’ (B56 or PR61), B’’ (PR48/PR72/PR130), or B’’’ (PR93/PR110)^29,31,32^. We have identified B55α as the cognate regulatory subunit of PP2A, and the enzymatic activity of PP2A holoenzyme counteracts PINK1-initiated mitophagy.

## Results

### PP2A is the phosphatase for pUb

We prepared pUb as previously described^8^, and collected two-dimensional ^1^H-^15^N heteronuclear single-quantum coherence (HSQC) NMR spectra. In the presence of cell lysate at a 1:1 (w/w) ratio for 4 h at room temperature, the HSQC spectrum of pUb reverts to that of unmodified ubiquitin (Supplementary Fig. 1). Note that the spectral changes were not due to protein degradation, since all the peaks can be readily assigned (Supplementary Fig. 1b). Therefore, the cell lysate has phosphatase activity specific for the removal of S65 phosphoryl group from pUb.

To monitor the dephosphorylation of pUb in real-time, we conjugated a trifluoromethyl probe at the K48C site of ubiquitin^33,34^, and collected a series of one-dimensional ^19^F NMR spectra. As shown previously, pUb fluctuates between two conformational states, namely the relaxed state and retracted state^8,9,13^. The two conformational states are manifested as two distinct ^19^F peaks at -83.70 ppm and -83.86 ppm, respectively (Supplementary Fig. 2a)^33^. The addition of cell lysate causes a gradual disappearance of these two ^19^F peaks, while at the same time, a new peak appears at -83.52 ppm that corresponds to the unmodified ubiquitin. Phosphatases can be inhibited by small molecules in accordance with their catalytic mechanism^35-37^. The addition of okadaic acid or cantharidin inhibits the phosphatase activity of the cell lysate and slows the pUb dephosphorylation rate by more than fourfold (Supplementary Fig. 2). Okadaic acid and cantharidin are inhibitors of PP2A and PP1^35,38,39^, and therefore, our ^19^F-NMR screening narrowed the list of the pUb phosphatase.

To further identity the phosphatase for pUb, we prepared PP2A and PP1 core enzymes recombinantly and titrated them to ^19^F-labeled pUb. The addition of 0.5 µM PP2A rapidly eliminates the two pUb peaks and expedites the concomitant appearance of the ^19^F peak corresponding to the unmodified ubiquitin (Supplementary Fig. 3). The initial dephosphorylation rate is estimated at ∼1.6 µM/min (Fig. 1a). In comparison, the addition of 0.5 µM PP1 core enzyme causes little change to the ^19^F peak intensities, comparable to those of untreated pUb. Further assessment of the HSQC spectra of PP2A-treated pUb also indicates a rapid reversal to that of unmodified ubiquitin (Fig. 1b). Together, PP2A is a *bona fide* phosphatase of pUb.

**Fig. 1.**
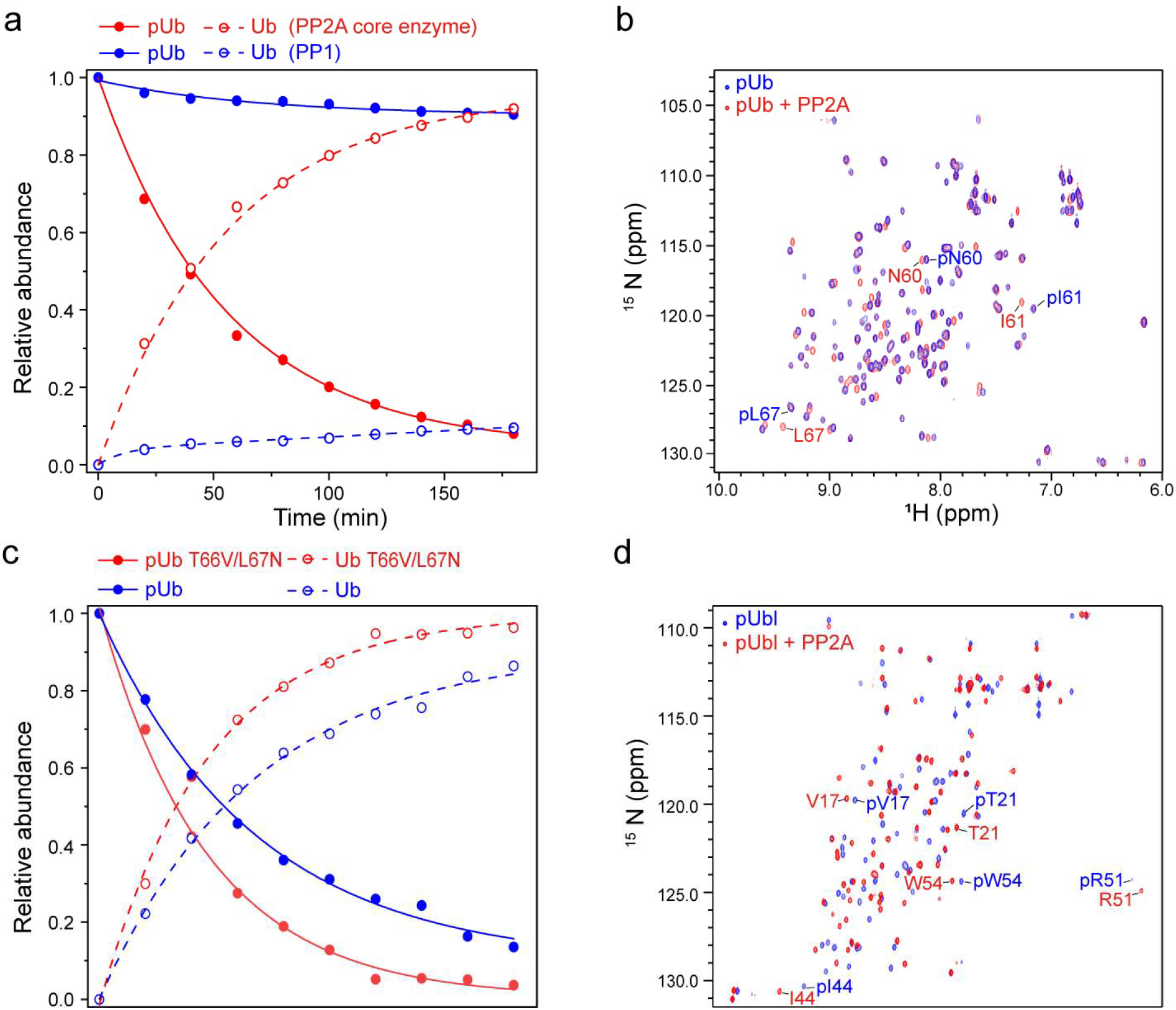
PP2A is the phosphatase for pUb and Parkin pUbl domain. **a,c** Normalized to the ^19^F peak intensities measured in real-time, referencing to the intensities at time 0; the Ub peak intensity was calculated as the total intensity subtracting remaining pUb peak intensity. 100 µM phosphorylated substrate (either ^19^F-probe conjugated ubiquitin or ubiquitin mutant) was mixed with 0.5 µM enzyme, and the reaction was allowed to proceed for 0-180 min (c.f. Supplementary Fig. 3 and Fig. 4). **b,d** Overlay of the 2D ^1^H-^15^N HSQC spectra of pUb and isolated pUbl (residues 1-76) from Parkin in the presence of purified PP2A core enzyme. 100 µM phosphorylated substrate (either ^15^N-labeled ubiquitin or Ubl from Parkin) was mixed with 0.5 µM enzyme, and the reaction was allowed to proceed for 1 h. Well-isolated peaks for the phosphorylated and unphosphorylated proteins are labeled.

### PP2A is the phosphatase for pParkin

PINK1 phosphorylates Parkin at its N-terminal Ubl domain, a critical step in the initiation of mitophagy^10,11^. Here we discovered that PP2A also dephosphorylates pUbl—the addition of 0.5 µM of the core enzyme reverts the NMR spectrum to that of the unphosphorylated Ubl (Fig. 1d). Significantly, the dephosphorylation of the isolated pUbl from Parkin by PP2A is much faster than the dephosphorylation of pUb, as the pUbl band completely disappears in less than 15 min with the addition of PP2A core enzyme (Fig. 2b). In comparison, the pUb level drops to ∼50% after >30 min incubation with PP2A, as shown with both Phos-tag gel and ^19^F NMR analyses (Fig. 2a and Fig. 1c).

**Fig. 2.**
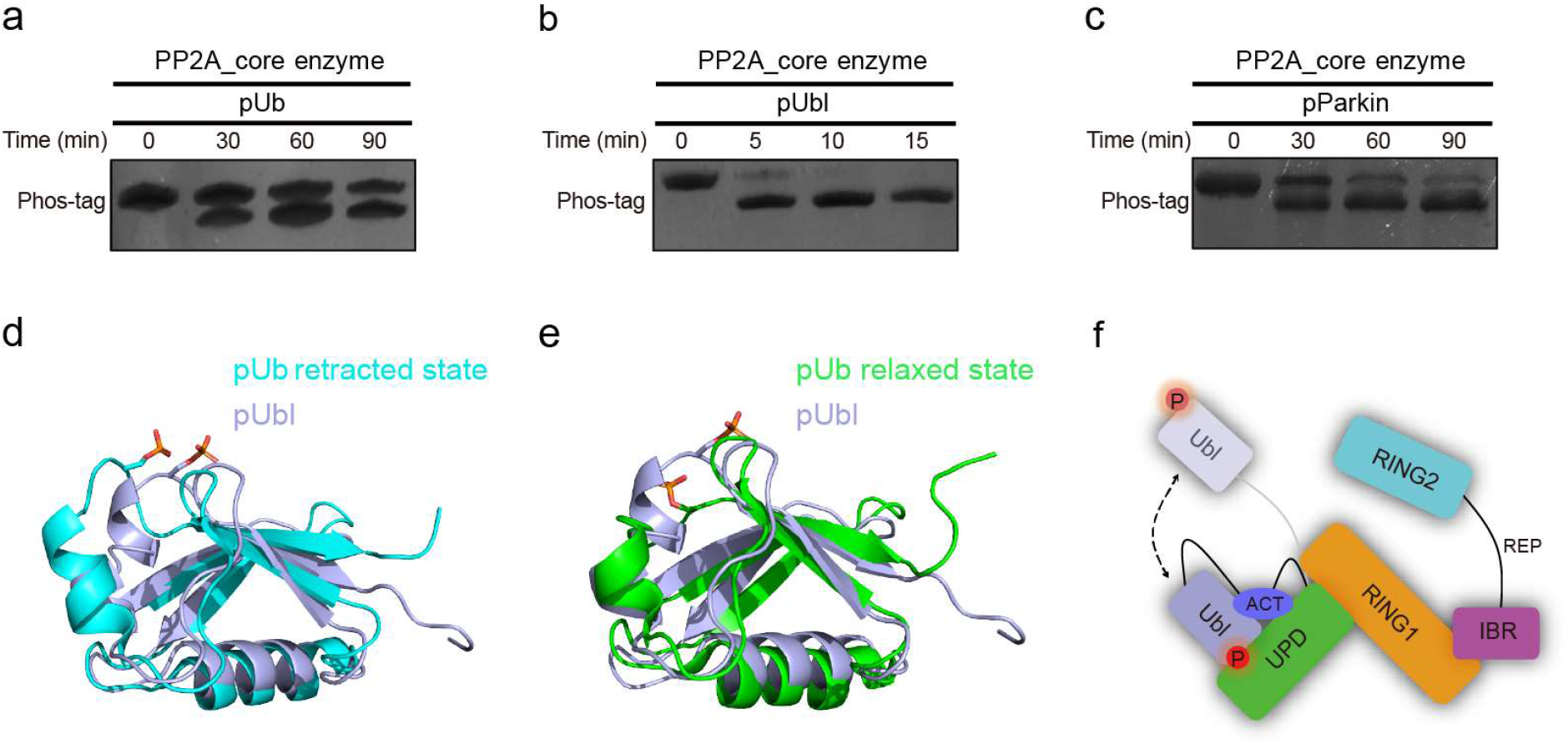
Substrate preference of PP2A. **a,b** Phos-tag gel analysis indicates that Parkin pUbl is a better substrate of PP2A core enzyme than pUb. **c** Phos-tag gel indicates that the dephosphorylation of pUbl by PP2A core enzyme is slower when the pUbl domain is part of the full-length Parkin. **d,e** Superposition of the structures of Parkin pUbl domain (PDB accession code 5TR5) with pUb retracted state (PDB accession code 5XK4), and with pUb relaxed state (PDB accession code 5XK5), affording a root-mean-square deviation of 1.44 Å and 1.64 Å, respectively, for the backbone heavy atoms. **f** Schematic illustrating the partial or transient protection of the phosphorylation site of Parkin pUbl domain.

The difference in PP2A dephosphorylation rates has a structural explanation. Residue S65 in Parkin is part of an extended loop, whereas residue S65 in ubiquitin only becomes exposed in the retracted state^8,13^. Indeed, the Parkin Ubl domain is more structurally similar to the pUb retracted state than to the relaxed state, and its S65 phosphoryl group faces outwards and is readily accessible. (Fig. 2d,e). Previously Komander and coworkers designed a ubiquitin double-mutant and drove pUb completely to the retracted state^40^. We found that PP2A dephosphorylated this mutant pUb more efficiently than the wild-type pUb (Supplementary Fig. 4 and Fig. 1c), indicating that PP2A preferentially catalyzes pUb in its retracted state. Moreover, pUb only partially samples the retracted state^8,13^, which further accounts for the slower dephosphorylation rate of pUb than pUbl of Parkin.

The binding of pUb to Parkin allosterically triggers the release of the Ubl domain that can be subsequently phosphorylated by PINK1^10-12^. Nevertheless, we found that PP2A dephosphorylated pUbl domain as part of the full-length Parkin less efficiently than an isolated pUbl, with about a tenfold reduction in the phosphorylation rate (Fig. 2c). The difference arises from the lower accessibility of the substrate phosphoryl group in the full-length Parkin, as the pUbl domain would competitively bind to the UPD domain with the S65 phosphoryl group largely occluded (Fig. 2f)^11^.

### B55α is the regulatory subunit for pUb and pParkin dephosphorylation

A PP2A holoenzyme contains a regulatory subunit that confers subcellular localization and substrate specificity^31,32^. Using FLAG-tagged pUb and GST-tagged PP2A scaffolding subunit A, we pooled cell lysate, performed mild cross-linking reactions, and pulled down pUb-interacting proteins with consecutive use of FLAG and GST affinity resins. In this way, we identified regulatory subunit B55α in the mixture using high-resolution mass spectrometry (Supplementary Fig. 5).

We prepared the B55α recombinantly and mixed it with the PP2A core enzyme at a 1:1 ratio. The addition of the regulatory subunit accelerates pUb dephosphorylation, as assessed from the changes of ^19^F NMR peak intensities (Supplementary Fig. 6). PP2A holoenzyme also catalyzes the dephosphorylation of pParkin more efficiently than the PP2A core enzyme, as assessed with the Phos-tag gel (Fig. 3a,b). Analysis using mass spectrometry also revealed more rapid appearance of 51,796 Da peak, corresponding to the unmodified Parkin, upon the treatment of PP2A holoenzyme than upon the treatment of PP2A core enzyme added at the same molar ratio for the same duration of incubation time (Fig. 3c,d). As such, B55α together with the core enzyme of PP2A makes up the PP2A holoenzyme and dephosphorylates both pUb and pParkin.

**Fig. 3.**
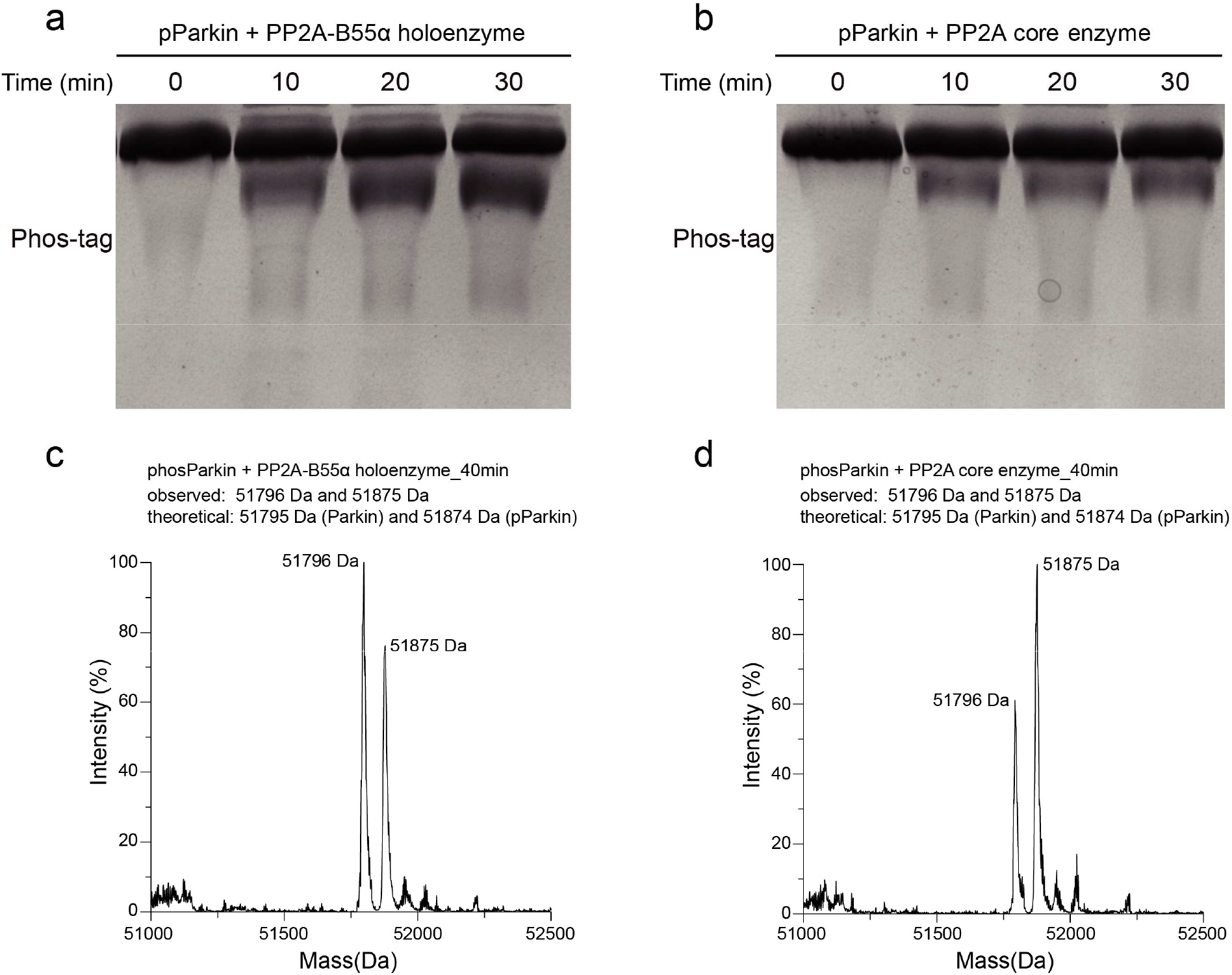
B55α accelerates PP2A dephosphorylation of pParkin. **a,b** In the presence of regulator subunit B55α, PP2A holoenzyme dephosphorylates pParkin faster than PP2A core enzyme (comprising only catalytic Cα and scaffolding Aα subunits). **c,d** Mass spectrometry analysis of pParkin dephosphorylation by PP2A holoenzyme and core enzyme, respectively. The reactions were performed for 40 min at room temperature. The dephosphorylation causes a mass decrease of 79 Da. Note that Parkin has two additional residues, proline and glycine, at the N-terminus, after PreScission protease cleavage to remove the GST tag.

### PP2A modulates pUb and pParkin levels in cells

Administration of CCCP or oligomycin and antimycin A (O/A) can lead to the loss of mitochondrial membrane potential and trigger PINK1-Parkin-mediated mitophagy^5,41^. Accordingly, increased phosphorylation levels of Parkin and ubiquitin were observed (Fig. 4). Note that the phosphorylated polyubiquitin has a distinct ladder from that of the polyubiquitin (Fig. 4c), suggesting PINK1’s selectivity for ubiquitin linkage and length^13^. Transient transfection of the PP2A catalytic subunit causes a slight increase of the protein band at ∼36 kDa. Consequently, the pParkin and pUb protein bands, otherwise strongly elicited by CCCP and O/A treatment, have decreased in intensities (Fig. 4a,c, 5^th^, and 6^th^ lanes).

**Fig. 4.**
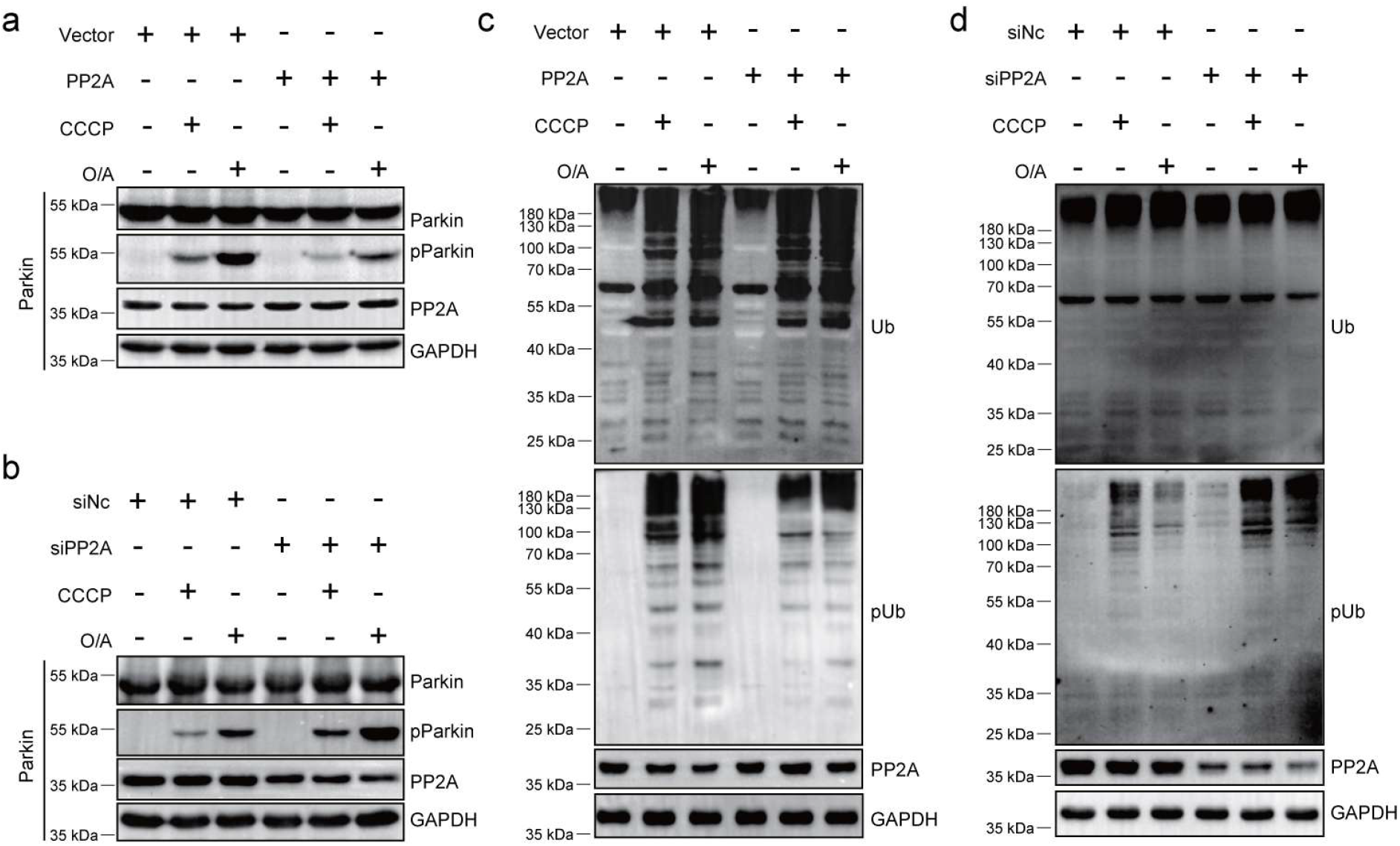
PP2A modulates phosphorylation levels of pParkin and pUb. **a,c** Transient transfection of PP2A catalytic subunit causes an increase of the phosphatase level, and a decrease of pParkin and pUb levels, respectively. **b,d** Knockdown of the phosphatase level with siRNA specific for PP2A catalytic subunit causes an increase of pParkin and pUb levels, respectively. Note that the overall Parkin and polyubiquitin levels remain unchanged, regardless of the changes of the PP2A level. The increase of pParkin and pUb levels were first elicited with the treatment of CCCP (10 µM for 4 h) or oligomycin/antimycin A (both at 250 nM for 2 h).

Conversely, introducing a siRNA specific for PP2A catalytic subunit lowers the protein band intensity at 36 kDa. As a result, the pParkin and pUb protein levels increase, indicating a cumulative effect with the CCCP and O/A treatment (Fig. 4b,d, 5^th^, and 6^th^ lanes). Note that PP2A is highly abundant in the cell, yet the protein level still changes upon transient transfection or siRNA treatment. Importantly, as a control, regardless of the PP2A protein level, the overall Parkin and ubiquitin levels remain unchanged. Taken together, PP2A actively dephosphorylates pParkin and pUb in cells and counteracts PINK1 kinase activity.

### PP2A inhibits Parkin translocation to mitochondria

Parkin recruitment to mitochondria is a key step in the initiation of mitophagy, as the activated Parkin ubiquitylates a multitude of OMM proteins^19,42^. We used HeLa cells stably transfected with YFP-tagged Parkin^42^ and treated the cells with CCCP or O/A. Translocation of Parkin, as characterized by the colocalization of YFP fluorescence with mitochondria marker TOM20, could be observed (Fig. 5a, left panel), which increased over time. However, the transient transfection of PP2A catalytic subunit causes a significant reduction of Parkin colocalization to the mitochondria at different durations of CCCP or O/A treatment (Fig. 5a, right panel, and Fig. 5b).

**Fig. 5.**
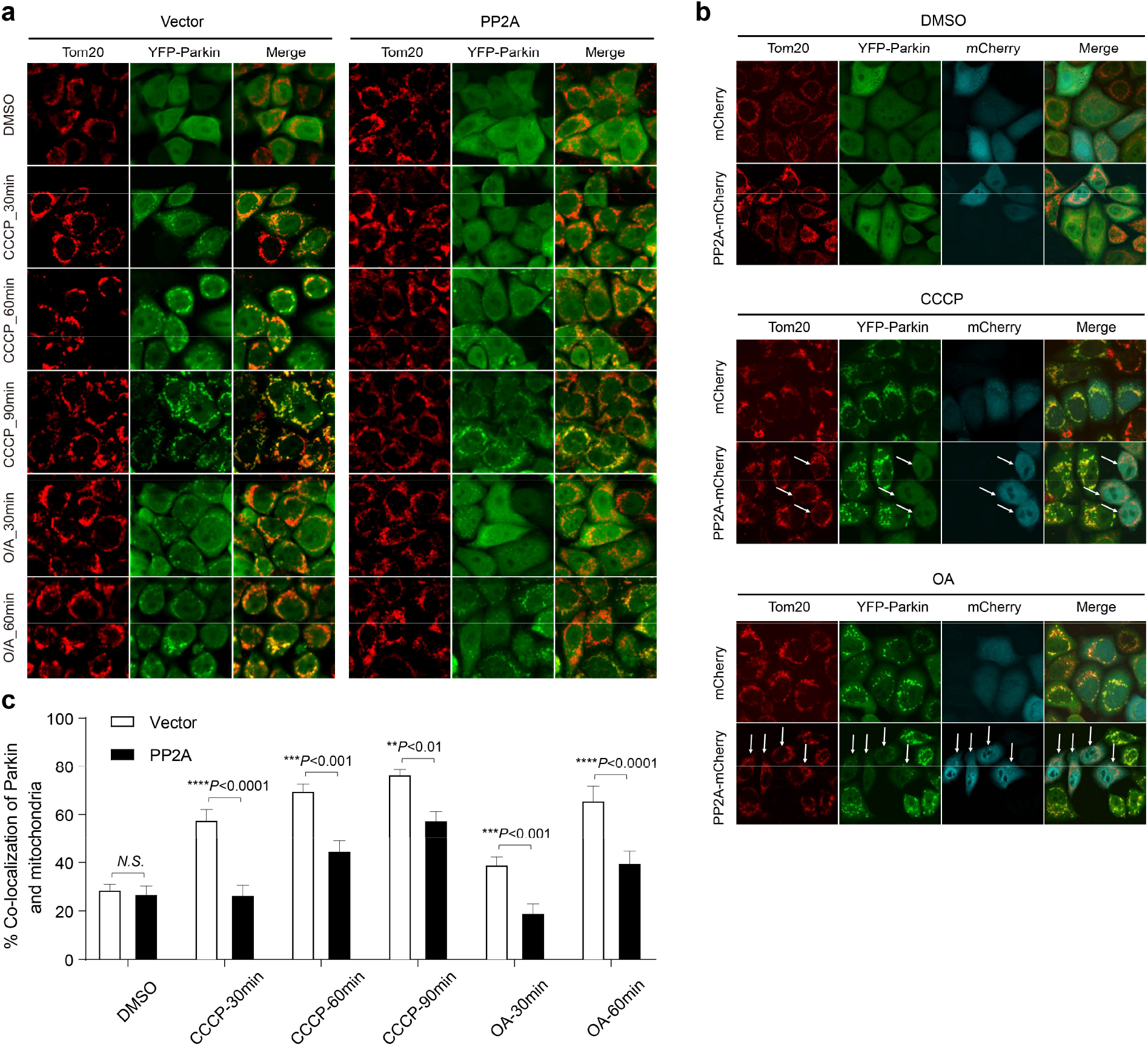
PP2A dephosphorylation inhibits Parkin translocation to mitochondria. **a** Transient transfection of PP2A (core enzyme of Aα and Cα subunits) prevents Parkin’s localization at depolarized mitochondria. Hela cells stably transfected with YFP-Parkin (green) show an increased mitochondrial localization upon the treatment with CCCP (10 µM for 0-90 min) or O/A (both at 250 nM for 0-60 min). DMSO was used as a control; TOM20 (red) immunofluorescence staining delineates mitochondria. **b** Transient transfection of YFP-Parkin HeLa cells with mCherry or PP2A-Cα-mCherry. After the CCCP (10 µM for 2 h) or O/A (both at 250 nM for 1 h) treatment, YFP-Parkin puncta were observed for cells without mCherry fluorescence (cyan) in PP2A-Cα-mCherry transfected cells. **c** Quantitative analysis for mitochondria localization of Parkin in (**a**). Quantitation was averaged from three independent experiments. All data were analyzed by two-way analysis of variance (ANOVA). Data are displayed as mean ± SEM. Statistical significance is displayed as \*\**P* < 0.01, \*\*\**P* < 0.001 and \*\*\*\**P* < 0.0001, *N*.*S*.: not significant.

To further explore the over-expression of PP2A on Parkin translocation, we fused a mCherry gene at the 3’-end of PP2A catalytic subunit Cα, and assessed the mCherry fluorescence in the cells upon transient transfection. PP2A was found evenly distributed in the cells, while transient transfection of PP2A-mCherry did not affect Parkin translocation (Fig. 5b). Upon CCCP or O/A treatment, Parkin translocation to mitochondria, manifested by the colocalization of YFP fluorescence and TOM20 staining. Significantly, Parkin translocation could only be observed in mCherry-negative cells (Fig. 5b), whereas cells expressing PP2A-Cα-mCherry showed little change in YFP-Parkin fluorescence in response to CCCP or O/A treatment. As such, PP2A completely inhibits the translocation and recruitment of Parkin to mitochondria.

The absence of Parkin translocation can be due to several reasons. First, as the OMM pUb is dephosphorylated by PP2A, Parkin can no longer be recruited to mitochondria. On the other hand, in comparison to pUb, pParkin is more efficiently dephosphorylated by PP2A, either at mitochondria or in the cytoplasm. Consequently, the majority of Parkin remains in the cytoplasm.

### PP2A negatively regulates mitophagy

Having established that PP2A inhibits Parkin translocation to mitochondria, we further investigated the role of PP2A in regulating mitophagy. We used Mito-QC to assess the progress of mitophagy, as the fusion between mitochondrion and lysosome leads to distinctive red punctum due to a decrease in the pH^43^. The administration of CCCP or O/A increased the number of mito-lysosomes in the control experiments, visible by Mito-QC (Fig. 6). Transient transfection and overexpression of PP2A Cα subunit, however, inhibits this process, with the mito-lysosomes remaining at a relatively low level (Fig. 6a,b). On the other hand, the introduction of siRNA specific for PP2A decreased the phosphatase level in the cell (Fig. 4b). Consequently, a greater number of mito-lysosomes was observed upon CCCP or O/A treatment (Fig. 6c,d). As such, PP2A dephosphorylation of pUb and pParkin forestalls mitophagy.

**Fig. 6.**
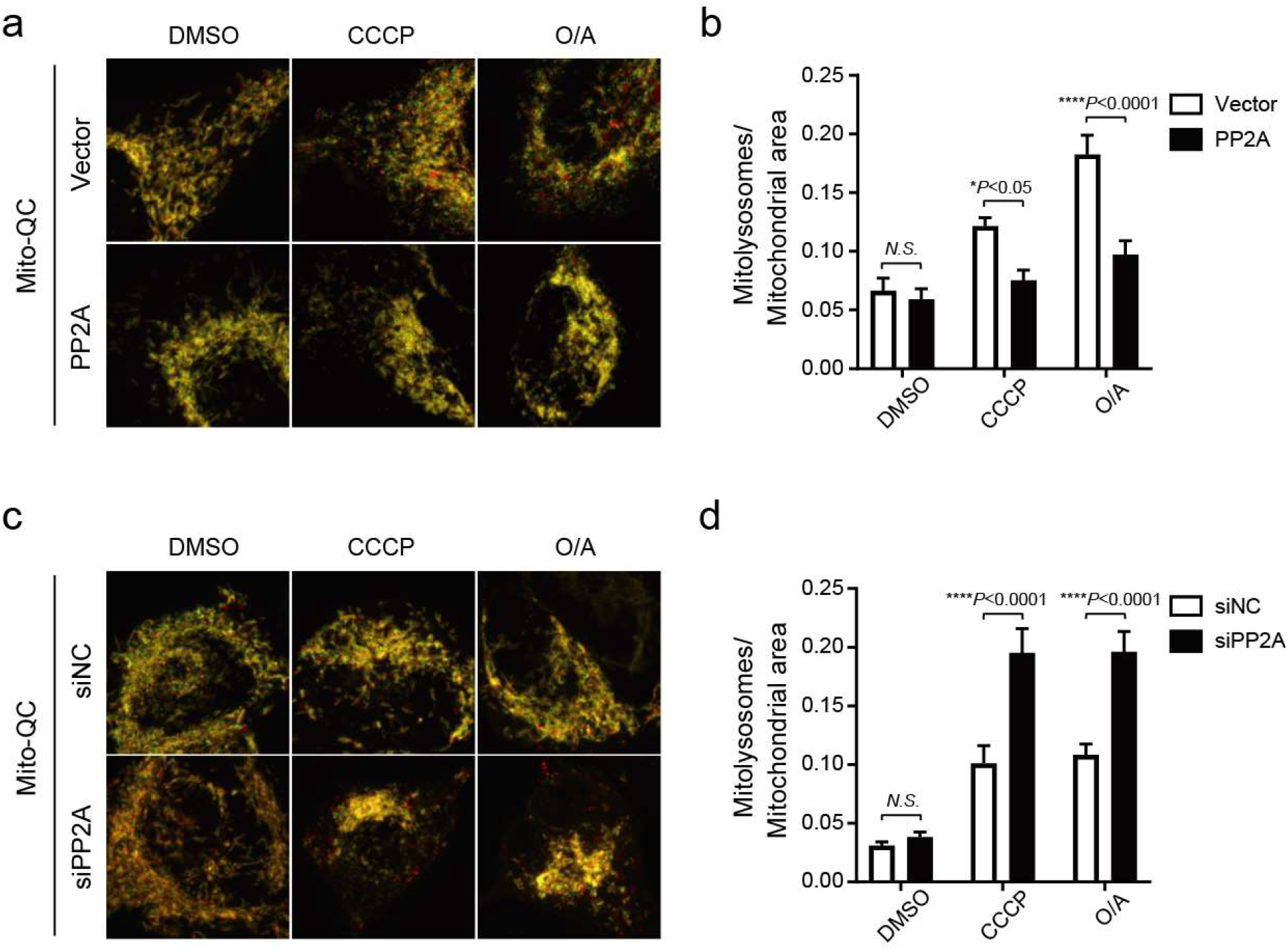
PP2A modulates the extent of mitophagy. Mitophagy is assessed with Mito-QC, as each red punctum indicates the fusion between mitochondrion and lysosome. **a, c** Transient transfection of SH-SY5Y cells with the plasmid expressing PP2A Cα subunit, and siRNA specific for PP2A Cα, decreases and increases the number of visible red puncta, respectively. **b,d** Quantitative analysis of the extent of mitophagy from the Mito-QC assay shown in (**a**) and (**c**), respectively. CCCP (10 µM for 4 h) or O/A (both at 250 nM for 2 h) were administrated to the cells to depolarize mitochondrial membrane potential and trigger mitophagy. The ratio of the number of mito-lysosomes to the mitochondrial area was quantitated and averaged over three independent experiments. Overall data are represented as mean ± SEM. Statistical significance is displayed as \**P* < 0.05, and \*\*\*\**P* < 0.0001 (two-way ANOVA), *N*.*S*.: not significant.

## Discussion

Mitophagy maintains the integrity of the mitochondrial network and promotes cellular homeostasis and survival. Yet aberrant and overactivated mitophagy is deleterious, as cells can be left with an insufficient quantity of mitochondria for energy production and, at the same time, an excessive number of lysosomes^44,45^. Indeed, depending on the stress conditions, mitochondria may only isolate the damaged portion and shed vesicles without the engulfment of the entire organelle^46,47^. Therefore, mitophagy has to be tightly regulated with both initiating and braking mechanisms in place.

Over the past decade, a lot has been learned about mitophagy initiation, in which kinase PINK1 and E3 ligase Parkin play essential roles^48,49^. In comparison, much less is known about the mitophagy braking system. Previously, two enzymes were identified as the phosphatases for pUb^24,25^. In one study, the authors proposed that ubiquitin dephosphorylation indirectly impairs Parkin mitochondrial translocation, but fell short in providing direct evidence for pParkin dephosphorylation^26^. In the present study, based on NMR, biochemical and imaging results, we unambiguously identified PP2A as the phosphatase specific for pParkin. The heterotrimeric protein phosphatase PP2A that includes the catalytic Cα subunit, the scaffolding Aα subunit, and regulatory B55α subunit, dephosphorylates both pUb and pParkin, blocks Parkin mitochondrial translocation, and inhibits mitophagy.

PINK1-phosphorylated ubiquitin participates in a multitude of cellular events, some of which are unrelated to mitophagy^13,27,28,50^. In comparison, Parkin is directly responsible for the ubiquitinylation of downstream OMM proteins, and activated Parkin has evolved poor specificity over substrate proteins but high spatial selectivity at the depolarized and damaged mitochondria^51^. Therefore, pParkin dephosphorylation is more direct and economical to forestall mitophagy. On the other hand, the dephosphorylation of ubiquitin may prevent Parkin mitochondrial translocation and activation in the first place (Fig. 7).

**Fig. 7.**
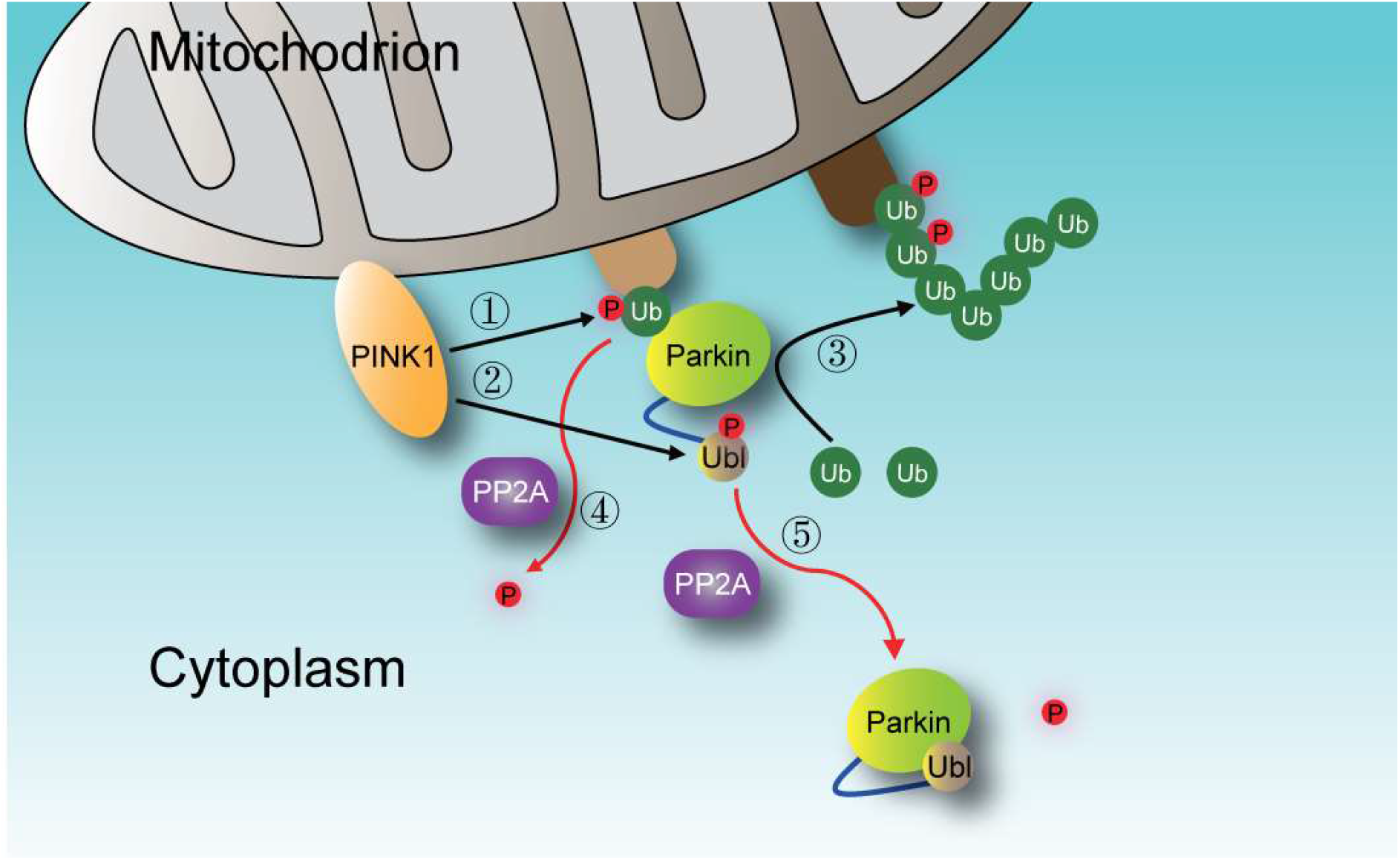
A scheme explaining how PP2A forestalls mitophagy. Parkin, an E3 ligase that ubiquitylates a multitude of OMM proteins, functioning as a master regulator of mitophagy through a feedforward mechanism. Upon mitochondria damage and membrane potential depolarization, PINK1 is activated and phosphorylates ubiquitin➀. The pUb recruits Parkin to mitochondria, and concomitantly, allosterically unhinge Parkin N-terminal Ubl domain, which can be readily phosphorylated by PINK1 ➁. The phosphorylated Parkin has full E3 ligase activity and ubiquitinylates mitophagy-related proteins➂. In a reversal, PP2A dephosphorylates both pUb➃and pParkin. Consequently, Parkin reverts back to the inactive auto-inhibited conformational state and is released back to the cytoplasm➄.

Previous studies have identified deubiquitinases responsible for removing polyubiquitin modifications from Parkin substrate proteins and inhibiting mitophagy^20-22^. However, phosphorylated polyubiquitins have increased conformational heterogeneities^8,13^, and are resistant to the hydrolysis of deubiquitinases^13^. Thus, the dephosphorylation of pUb has to occur first, before deubiquitinases can actively hydrolyze polyubiquitins and exert a negative regulatory effect on mitophagy. In this regard, inhibiting the activity of E3 ligase Parkin by dephosphorylating pParkin is more efficient by preventing the ubiquitylation of OMM proteins.

PINK1-Parkin signaling is known involved in the pathogenesis of Parkinson’s disease and many other neurodegenerative diseases^45,52^. On the other hand, PP2A is highly expressed in the brain^53^, and its dysregulation has also been implicated in the onset of Parkinson’s disease as well^54,55^. In particular, PP2A with B55α regulatory subunit has been shown responsible for dephosphorylating α-synuclein and tau proteins^56,57^. As a result, PP2A has been recognized as an attractive drug target for the treatment of cancer, inflammatory and neurodegenerative diseases^58^. Our present finding adds two important substrate proteins for PP2A, and demonstrates the functional relevance of PP2A in Parkin-mediated mitophagy.

## Methods

### Protein sample preparation

K48C, T66V/L67N, and N-terminal FLAG mutations were introduced to the human ubiquitin gene using the QuikChange method, and the resulting genes were cloned into a pET11a vector. Except for FLAG-Ub, the expression and purification of the ubiquitin proteins followed the established protocol^8^. As for FLAG-Ub, the protein was purified with Source-Q (GE Healthcare) and Sephacryl S100 columns (GE Healthcare) in tandem. For uniform ^15^N-labeling, BL21* cells transfected with ubiquitin plasmid were grown in M9-minimal medium with ^15^NH_4_Cl (Cambridge Isotope Laboratory) as the sole nitrogen source.

PP1α (Uniprot ID P62136) was cloned to a pET-11a vector with an 8xHis tag appended at its N-terminus. BL21* cells were used to express proteins. Cells were grown in LB medium at 37 °C and induced by 0.2 mM IPTG for 20 hours at 18 °C before harvested. The protein was purified with Ni-NTA Superflow affinity column (GE Healthcare) and Sephacryl S100 columns (GE Healthcare) in tandem.

PP2A scaffolding subunit Aα (Uniprot ID P30153) and regulatory subunit B55α (Uniprot ID P63151) were cloned to the pGEX-6P-1vector, respectively. BL21* cells were used to express proteins. Cells were grown in LB medium at 37 °C and induced by 0.2 mM IPTG for 20 hours at 18 °C before harvested. The proteins were purified using Glutathione Sepharose 4B (GE Healthcare) and Sephacryl S100 columns (GE Healthcare) in tandem.

PP2A catalytic subunit Cα (Uniprot ID P67775) was cloned into the pFastBac1 vector with a 10xHis tag appended at the N terminus and was expressed in Sf9 cells (Gibco, catalog number 12659017). The cells were cultured at 27 °C for 60 h after being infected with baculoviruses using the Bac-to-Bac system (Life Technologies). The recombinant proteins were purified using a Ni-NTA Superflow affinity column (GE Healthcare) and Source-Q (GE Healthcare) in tandem.

Parkin (Uniprot ID O60260) was cloned to a pET11a vector, with a His-SUMO tag appended at the N-terminus. BL21* cells were used to express proteins. Cells were grown in LB medium at 37 °C and induced by 0.1 mM IPTG and 200 mM ZnCl_2_ for 20 hours at 18 °C before harvested. The protein was purified with the Ni-NTA Superflow affinity column (GE Healthcare) and Sephacryl S100 column (GE Healthcare) in tandem. The isolated Parkin Ubl domain (residues 1-76 from the full-length human Parkin) was cloned to a pGEX-6P-1 vector. The protein was purified with Glutathione Sepharose 4B column (GE Healthcare) and Sephacryl S100 column (GE Healthcare) in tandem. The His-Sumo and GST tags at the N-terminus of the desired proteins were removed by incubation with PreScission Protease (Beyotime Biotechnology) at room temperature for 3 h.

PINK1 from body louse (phPINK1, Uniprot ID E0W1I1) was prepared as previously described^8^. To phosphorylate ubiquitin and Parkin, PINK1, ubiquitin (or Parkin), MgCl_2_, and ATP were mixed at the molar ratio of 1:100:5,000:5,000 in 20 mM Tris•HCl buffer (pH 8.0, containing 1 mM DTT). The reaction proceeded overnight at room temperature. To phosphorylate the isolated Parkin Ubl domain, the reaction was carried out with the same molar ratio and conditions, but only for 30 min. The product proteins pUb, pParkin, and pUbl were purified on Source-Q column (GE Healthcare), buffer-exchanged; the modifications were confirmed with Vion IMS QTOF Mass Spectrometer (Waters).

3-Bromo-1,1,1-trifluoroacetone (BFA) (J&K Scientific, catalog number 312226) were dissolved in DMSO to 5 M concentration as a stock solution. The K48C mutant of ubiquitin was desalted (HiPrep 26/10 Desalting column, GE Healthcare), and mixed with BFA at a molar ratio of 1:10 in 20 mM pH 7.4 HEPES buffer. The conjugation reaction was performed at room temperature for 2 h. The product was further purified using the desalting column, and the modification was confirmed with Vion IMS QTOF Mass Spectrometer (Waters).

For plasmids used in cell experiments, Parkin and PP2A-Cα-IRES-Aα were cloned to a pcDNA3.1 vector (IRES means the two genes are in the same vector but under separate promoters). PP2A-Cα alone was also cloned to a pcDNA3.1-mCherry vector.

### Phos-tag gel

To evaluate the phosphorylation level of the full-length Parkin, the protein samples were resolved on a 12% polyacrylamide gel containing 50 μM Phos-tag acrylamide (Wako Chemicals, catalog number AAL-107) and 100 μM MnCl2. Tricine protein gel with 12% polyacrylamide containing 100 μM Phos-tag acrylamide and 200 μM MnCl_2_ was used to assess the phosphorylation level of isolated Parkin Ubl domain and Ub. For sample preparation, 100 μM pParkin, pUb, or pUbl was incubated in enzyme buffer (50 mM Tris, pH 7.4, 150 mM NaCl, 1mM MnCl_2_) with 0.5 μM PP2A (core enzyme or holoenzyme, as noted in the Results) or buffer only at the room temperature, with the incubation time noted in the Results.

### Mass spectrometry identification of PP2A regulatory subunit

To identify PP2A regulatory subunit, we prepared an excess amount of HeLa cell lysate by manual grinding in the lysis buffer (50 mM HEPES, pH 7.4, 150 mM NaCl) in the presence of protease inhibitors (Coolaber, catalog number SL1086), and mixed the lysate with recombinantly purified FLAG-pUb and GST-PP2A Aα subunit. Subsequently, 4 mM freshly prepared BS_3_ (ThermoFisher, catalog number 21580) was added. The cross-linking reaction was conducted at room temperature with gentle mixing and was quenched after 45 min with the addition of Tris-HCl buffer (pH 8.0) to a final concentration of 20 mM. The proteins were pooled onto GST affinity resin (GE Healthcare), incubated for 2 h at 4 °C, washed three times with the lysis buffer, and eluted with the addition of lysis buffer containing 20 mM L-Glutathione reduced (Aladdin, catalog number G105426). Eluted proteins were incubated with anti-Flag magnetic beads (Bimake, catalog number B26102) for 2 h at 4 °C. Subsequently, the beads were washed three times in lysis buffer, and the elution was done with the and eluted by addition of 0.3 mg/mL poly-FLAG peptide (Bimake, catalog number B23111) to the lysis buffer. The eluted proteins were subjected to trypsin digestion.

The resulting peptide mixture was subjected to an Easy-nLC 1200 nano HPLC (Thermo Scientific, San Jose, CA) coupled to a Q Exactive Orbitrap mass spectrometer (Thermo Scientific, San Jose, CA). Samples were resolved on a C18 RP-trap column (150 µm i.d. x 3 cm) and a C18 capillary column (150 µm i.d. x 15 cm) in-house packed column with ReproSil-Pur C18-AQ particles (1.9 µm, 120 Å). Mobile phase A has 0.1% FA in HPLC H_2_O, and mobile phase B has 80% acetonitrile, 0.1% FA and 20% HPLC H_2_O. The flow rate was 300 nL/min, and the elution gradient used for peptide separation was as the following: 5-10% buffer B for 1 min, 10-40% buffer B for 95 min, 40-60% buffer B for 8 min, 60-100% buffer B for 1 min, and finally at 100% buffer B for 15 min. The mass spectrometer was operated in data-dependent mode with one full MS scan at R = 70,000 (m/z = 200), followed by HCD MS/MS scans at *R* = 17,500 (m/z = 200), NCE = 30, with an isolation width of 1.8 m/z. The AGC targets for the MS_1_ and MS_2_ scans were 3e+6 and 1e+5, respectively; the maximum injection times for MS_1_ and MS_2_ were 50 ms and 100 ms, respectively. The pFind Studio 3 software^59^ was used to identify the interacting peptides. The search parameters were set as the following: Cys carbamidomethyl selected as a static modification and Met oxidation a variable modification, precursor mass tolerance of 20 ppm, fragment mass tolerance of 20 ppm, full tryptic cleavage constraints but allowing up to 3 missed cleavages, and a spectral false identification rate ≤1%.

### NMR Spectroscopy analysis

For ^1^H-^15^N HSQC NMR experiments, samples were prepared in 20 mM HEPES buffer (150 mM NaCl, pH 7.4, with 5% D_2_O), and NMR data were acquired at 298 K on a Bruker 600 MHz spectrometer equipped with a Z-gradient cryogenic probe. The data were processed using NMRPipe^60^ and analyzed using NMRViewJ^61^. The protein samples used in ^19^F NMR were prepared in 50 mM Tris buffer (150 mM NaCl, 1 mM MnCl_2,_ and pH 7.4, with 5% D_2_O). NMR experiments were recorded at 298 K on a 600 MHz spectrometer (Bruker) equipped with a H/F/(C, N) triple-resonance cryogenic probe. The ^19^F base frequency is 564.39 MHz, with the 0 ppm set for the peak from trichlorofluoromethane. The peak volumes in the one-dimensional ^19^F NMR spectra were obtained with Bruker TopSpin Version 3.5 and NMRViewJ^61^. Since the ^19^F peaks of the phosphorylated and unphosphorylated Ub T66V/L67N mutant could not be completely distinguished, we fitted the relative populations of the three peaks using the Lorentz function with OriginPro 2019 (OriginLab).

For the dephosphorylation reaction, HeLa cells were harvested, washed twice with ice-cold 1x PBS buffer, and lysed at 4°C in the RIPA buffer (Beyotime Biotechnology) with protease inhibitors (Coolaber, catalog number SL1086). Total protein concentrations were determined using a BCA assay kit (Beyotime Biotechnology). HeLa cell lysate was added to 100 µM ^15^N-labeled pUb at a 1:1 (w/w) ratio. The reaction proceeded at room temperature for 4 h. HeLa cell lysate was also added to 100 µM ^19^F-labeled pUb at a 1:1 (w/w) ratio for the dephosphorylation reaction at room temperature. To inhibit the dephosphorylation reaction, we added 1 μM okadaic acid (Aladdin, catalog number O275664) or 10 μM Cantharidin (Aladdin, catalog number C111020) to 100 μM ^19^F-labeled pUb, before mixing with the cell lysate at a 1:1 (w/w) ratio. A series of 1D ^19^F NMR data were collected at room temperature to monitor dephosphorylation reaction in real-time. In addition, we used recombinantly purified 0.5 μM PP2A (core enzyme or holoenzyme) or 0.5 μM PP1 core enzyme and added it into 100 μM ^19^F-labeled pUb for dephosphorylation.

### Cell Lines and treatment of the cells

HeLa cells stably expressing YFP-Parkin were kind gifts from Prof. Dajing Xia (Zhejiang University), as previously described^42^. The SH-SY5Y cells (CAS Center for Excellence in Molecular Cell Science, China) were seeded onto poly-D-lysine (Sigma-Aldrich, catalog number P7405) coated glass-bottom dishes (Cellvis, catalog number D29-20-1-N) for live-cell imaging. The SH-SY5Y cells, regular HeLa cells (CAS Center for Excellence in Molecular Cell Science, China), and YFP-Parkin HeLa cells were grown in Dulbecco’s modified essential medium (DMEM, Gibco, catalog number 11965092) supplemented with 10% heat-inactivated fetal bovine serum (Gibco, catalog number 2275120) at 37 °C, in the atmosphere of 5% CO_2_.

### Western blotting

Transient transfections of the plasmids were achieved with the use of Lipofectamine 2000 (Thermo Fisher Scientific, catalog number 11668019). When the cells reached a confluency of ∼90%, ∼4 µg plasmid of pcDNA3.1-PP2A-Cα-IRES-Aα or ∼2 µg plasmid of pcDNA3.1-PP2A-Cα-IRES-Aα with ∼2 µg plasmid of pcDNA3.1-Parkin were transfected into Hela cells; the control group was transfected with the same amount of empty vector. After 24 h culture at 37 °C, 10 µM CCCP (Sigma-Aldrich, catalog number C2759) or 250 nM/250 nM oligomycin (Abcam, catalog number ab141829) and antimycin A (Sigma-Aldrich, catalog number A8674) were added; the incubation durations were reported in the Results.

The transfection of siRNA was achieved with the use of Lipofectamine RNAiMAX (Thermo Fisher Scientific, catalog number 13778075) with a final concentration of 50 nM siRNA. The PP2A Cα siRNA and control siRNA have the sequences of 5’-GGAACUUGACGAUACUCUA-3’ and 5’-UUCUCCGAACGUGUCACGU-3’, respectively. When the cells reached a confluency of 60%, PP2A Cα siRNA or PP2A Cα siRNA with ∼2 µg pcDNA3.1-Parkin plasmid were transfected into Hela cells; the control group was transfected with control siRNA or control siRNA with ∼2 µg pcDNA3.1-Parkin plasmid. After 48-h culture at 37 °C, 10 µM CCCP or 250 nM/250 nM O/A were added.

Cells were washed 3 times with ice-cold 1X PBS buffer and lysed with the RIPA buffer (Beyotime Biotechnology) with protease (Coolaber, catalog number SL1086) and phosphatase inhibitors (Coolaber, catalog number SL1087) present. The cell lysate was quantified using the BCA protein assay kit (Beyotime Biotechnology) and was reduced and boiled before loading onto 12% polyacrylamide gel. The proteins were transferred from the gel to PVDF membranes. The membranes were blocked with 5% nonfat milk in TBS buffer containing 0.05% Tween-20 (TBST) for 1 h, incubated with primary antibodies in the blocking buffer (TBST buffer with 5% nonfat milk) overnight at 4°C, and then HRP-conjugated secondary antibodies (diluted 1:2000 in blocking buffer) for 2 h at room temperature. Protein quantity was extrapolated based on HRP activity using the Clarity™ Western ECL Substrate (Bio-Rad Laboratories), measured on a ChemiDoc XRS+ (BioRad), and analyzed using the Image Lab software (Bio-Rad Laboratories).

The primary antibodies used the Western blotting include those for Parkin (Cell Signaling Technology, catalog number 2132S, diluted at 1:1000 ratio), pParkin (Cell Signaling Technology, catalog number 36866S, diluted at 1:500 ratio), ubiquitin (Beyotime Biotechnology, catalog AF1705, diluted at 1:1000 ratio), pUb (Merck Millipore, catalog number ABS1513-I, diluted at 1:500 ratio), PP2A Cα (Thermo Fisher Scientific, catalog number MA5-32920, diluted at 1:2000 ratio), GAPDH (Proteintech, 60004-1-Ig, catalog number MA5-32920, diluted at 1:20000 ratio). Secondary antibodies were purchased from Beyotime Biotechnology, including HRP-conjugated goat anti-rabbit IgG (catalog number A0208) and HRP-conjugated goat anti-mouse IgG (catalog number A0216).

### Immunofluorescent Imaging

YFP-Parkin-HeLa cells were plated on glass coverslips pretreated with TC (Solarbio Life Sciences, catalog number YA0350). When the cell density reached 90%, ∼1 µg pcDNA3.1-PP2A-Cα-IRES-Aα or pcDNA3.1-PP2A-Cα-mCherry was transfected to YFP-Parkin-Hela cells; as a control, cells were transfected with ∼1 μg pcDNA3.1 or pcDNA3.1-mCherry plasmid. After 24 h culture at 37 °C, 10 µM CCCP or 250 nM/250 nM O/A were added.

Following CCCP or O/A treatment, we carefully aspirated the medium from the edge of the 24-well plate, and then added 500 µl 1xPBS buffer to rinse the cells. The cells were fixed with 4% paraformaldehyde diluted in 1x PBS for 15 min at 37°C. We then decanted paraformaldehyde solution, washed the cells three times with 500 µL 1xPBS, and added 200 µl 1xPBS containing 0.1% Triton X-100 for 20 minutes. The cells were then washed three times with 500 µL 1X PBS. 20 µL blocking buffer (1X PBS with 5% normal donkey serum; Yeasen Biotechnology, catalog number 36116ES03) was added to each cover slide for 30 min at room temperature. The primary antibody was diluted in the blocking solution at 1:200 ratio and was incubated overnight at 4 °C in a humidified box. The cells were washed three times with 1X PBS buffer, each time for ∼10 min. The secondary antibody was diluted at 1:200 certain ratio in the blocking buffer, and was applied for 2 h at room temperature in a humidified box protected from light. The cells were washed three times with 1x PBS buffer in the dark, with 10 minutes for each time. 5 µl DAPI Fluoromount-G™ (Yeasen Biotechnology, catalog number 36308ES11) was added to cover the cells. The slides were imaged under a Laser Scanning Confocal Microscope (Nikon, A1R-si) with a 60x immersion oil objective. The antibodies used include anti-Tom20 (Santa Cruz, catalog number SC-17764) and anti-Goat Mouse IgG (H+L) cross-adsorbed secondary antibody with Alexa Fluor 647 (Thermo Fisher Scientific, catalog number A-21235). The colocalized areas from YFP-Parkin and anti-Tom20 fluorescence were quantitated using Nikon NIS-Elements AR imaging software.

### Live-cell imaging and analysis of mitophagy

The residues 101–152 of human Fis1 cDNA were inserted into the BglII and EcoRI site of the pEGFP-C1 plasmid to construct the pEGFP-FIS1 plasmid. Then EGFP-Fis1 cDNA was amplified by PCR with the forward and reverse primers of primer 5′-TTAAGCTTCGATGGTGAGCAAGGGCGAGGAGCTGTT-3′ and 5′-TGGAATTCTCAGGATTTGGACTTGGACACAGCAAGTC-3′, respectively. The PCR product was inserted into the HindIII and EcoRI sites of the pmCherry-C1 plasmid to construct the Mito-QC plasmid^43^. SH-SY5Y cells were transfected with Mito-QC plasmid using Lipofectamine 3000 (Thermo Fisher Scientific, catalog number L3000015). Following the various treatments, cells were imaged at 37 °C using an LSM880 inverted confocal microscope (Zeiss) fitted with a ×63 oil objective at appropriate excitation (GFP at 488 nm and mCherry at 548 nm) and emission (GFP at 519 nm and mCherry at 562 nm) wavelengths. For quantification, red puncta in each cell were counted as mito-lysosomes, and the size of the correspondent cells was measured as mitochondrial area using ImageJ software (National Institutes of Health).

### Statistics and reproducibility

The data were processed with Prism 8.0 (GraphPad) and presented as means ±SEM. Quantitation of the colocalization and Mito-QC were based on three independent experiments, with 10-20 cells per combination of the treatment for colocalization analysis and 15-20 cells per combination for Mito-QC analysis. Statistical significance between multi-group comparisons was determined by two-way analysis of variance (ANOVA). Significance levels are indicated in the figure legends. *P* values are indicated with asterisks with \**P* < 0.05, \*\**P* < 0.01, \*\*\**P* < 0.001, \*\*\*\**P* < 0.0001, and *N*.*S*. for not significant.

## Acknowledgment

The work was supported by the National Key R&D Program of China (2018YFA0507700), the National Natural Science Foundation of China (81971304 to W-P.Z. and 31770799 to C.T.). The NMR experiments were performed at the National Center for Magnetic Resonance in Wuhan and the Beijing NMR Center and the NMR facility of the National Center for Protein Sciences at Peking University.

## Author contributions

S.Y. W.Z. and C.T. designed the experiments; S.Y. performed NMR and molecular biology experiments, with the assistance from Z.N. and L.M.; X.D. and Z.G. supported the collection and analysis of mass spectrometry and NMR data; X.L. and F.L. performed Mito-QC experiments under the supervision of W.Y.; S.Y., W.Z. and C.T. analyzed and interpreted the data; S.Y. and C.T. wrote the manuscript with inputs from all other authors.

## Additional information

### Competing interests

The authors declare no competing interests.

**Correspondence** and requests for materials should be addressed to Chun Tang.

---

**Supplementary Fig. 1.**
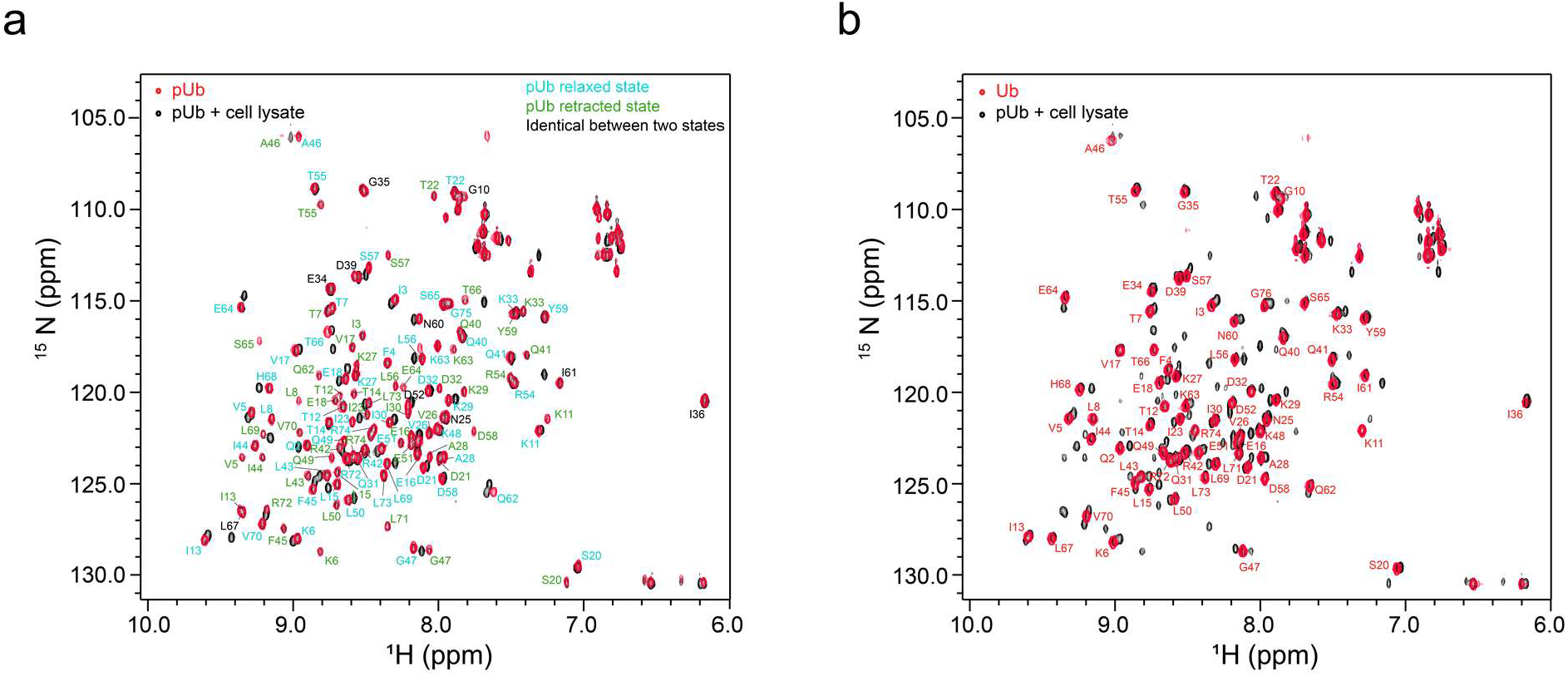
Overlay of ^1^H-^15^N HSQC spectra of ^15^N-labeled ubiquitin. **a** Comparison of the spectra of 100 µM pUb and pUb treated with HeLa cell lysate at a 1:1 (w/w) ratio for 4 h at room temperature. **b** Comparison of the spectra of pUb treated with cell lysate and the unmodified ubiquitin. The pUb peaks and unmodified Ub peaks are labeled. Note that 30-40% pUb remained after cell lysate treatment, judging by the relative peak intensities.

**Supplementary Fig. 2.**
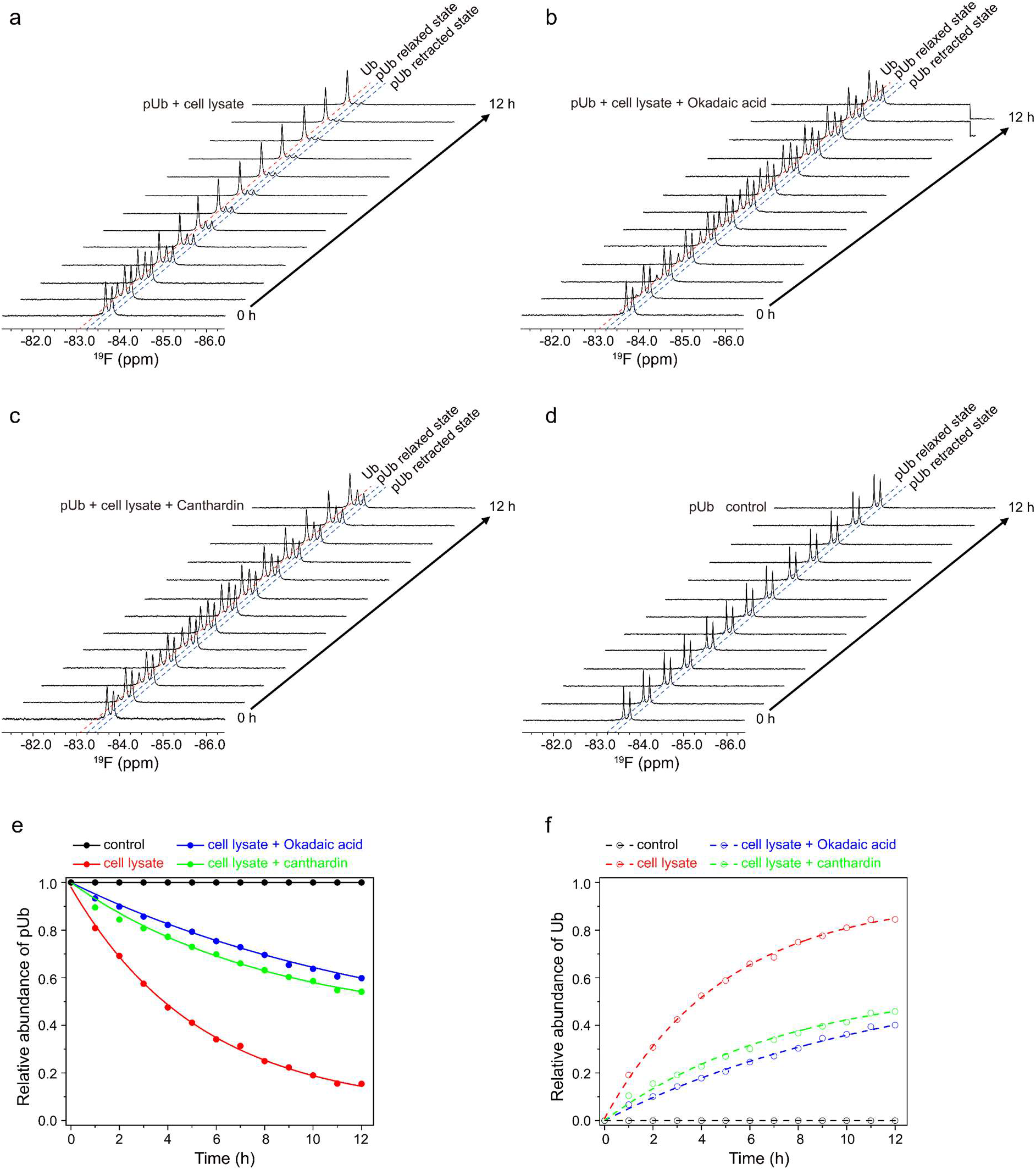
^19^F NMR real-time assessment of the inhibitory effect of small-molecule inhibitors. A trifluoromethyl probe was conjugated at the K48C site of ubiquitin. **a** The addition of HeLa cell lysate at a 1:1 (w/w) ratio to 100 µM ^19^F-labeled pUb causes gradual disappearance of the two pUb peaks and concomitant increase of the peak corresponding to the unmodified ubiquitin. **b,c** The addition of small-molecule inhibitors okadaic acid (1 µM) and cantharidin (10 µM), slows the relative changes of ^19^F peak intensities. **d** A stack of ^19^F NMR spectra of pUb alone, as a control without the addition of cell lysate and small-molecule inhibitors, showing no change in peak intensities over time. **e,f** Normalized to the ^19^F peak intensities, referencing to the intensities at time 0; the Ub peak intensity was calculated as the total intensity subtracting remaining pUb peak intensity.

**Supplementary Fig. 3.**
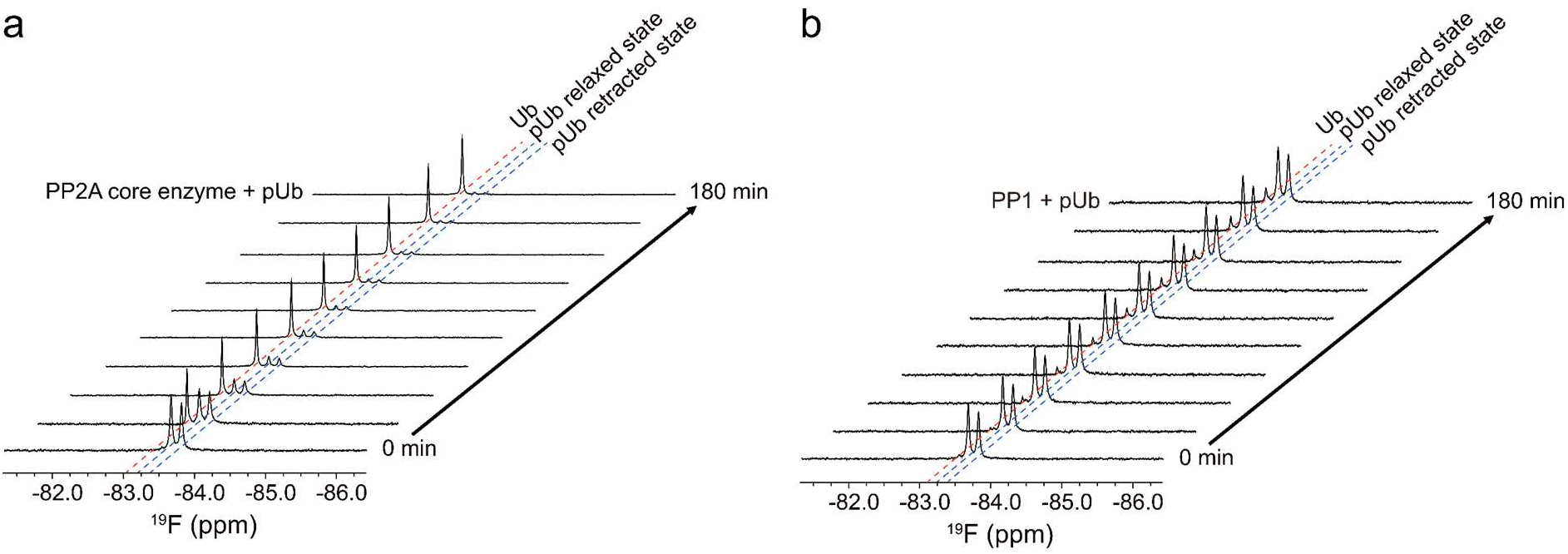
Real-time ^19^F NMR spectra of 100 µM pUb in the presence of purified phosphatases. **a** A stack of ^19^F NMR spectra in the presence of 0.5 µM PP2A core enzyme. **b** A stack of ^19^F NMR spectra in the presence of 0.5 µM PP1 core enzyme.

**Supplementary Fig. 4.**
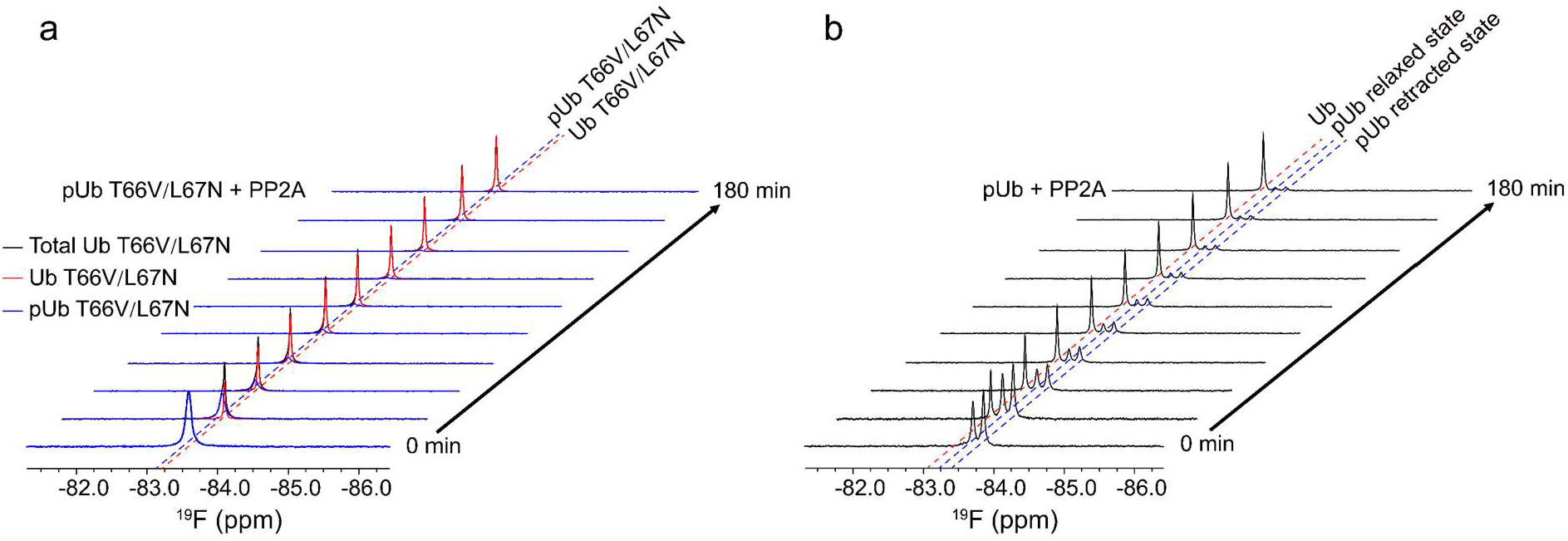
PP2A catalyzes pUb in its retracted-state conformation. **a** A stack of ^19^F NMR spectra monitoring the dephosphorylation process of phosphorylated T66/L67N ubiquitin double mutant. The mutations drove pUb completely to the retracted state. **b** A stack of ^19^F NMR spectra monitoring the dephosphorylation process of phosphorylated wild-type ubiquitin.

**Supplementary Fig. 5.**
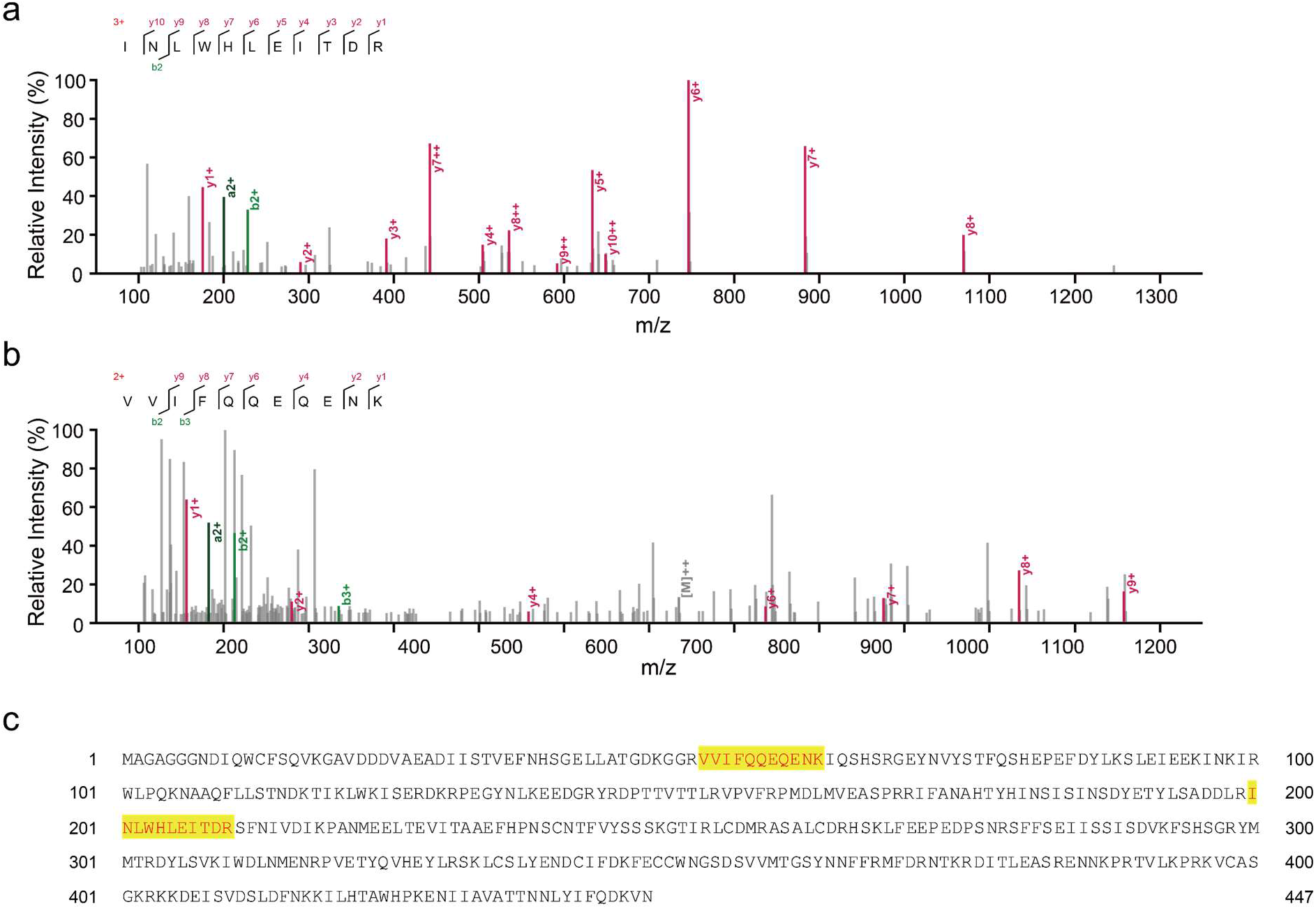
Identification of B55α as PP2A regulatory subunit. **a,b** High-resolution mass spectra of peptides identified from the cell lysate pulled down with GST-tagged PP2A scaffolding subunit and with FLAG-tagged pUb. **c** Matching these two peptides to the B55α sequence.

**Supplementary Fig. 6.**
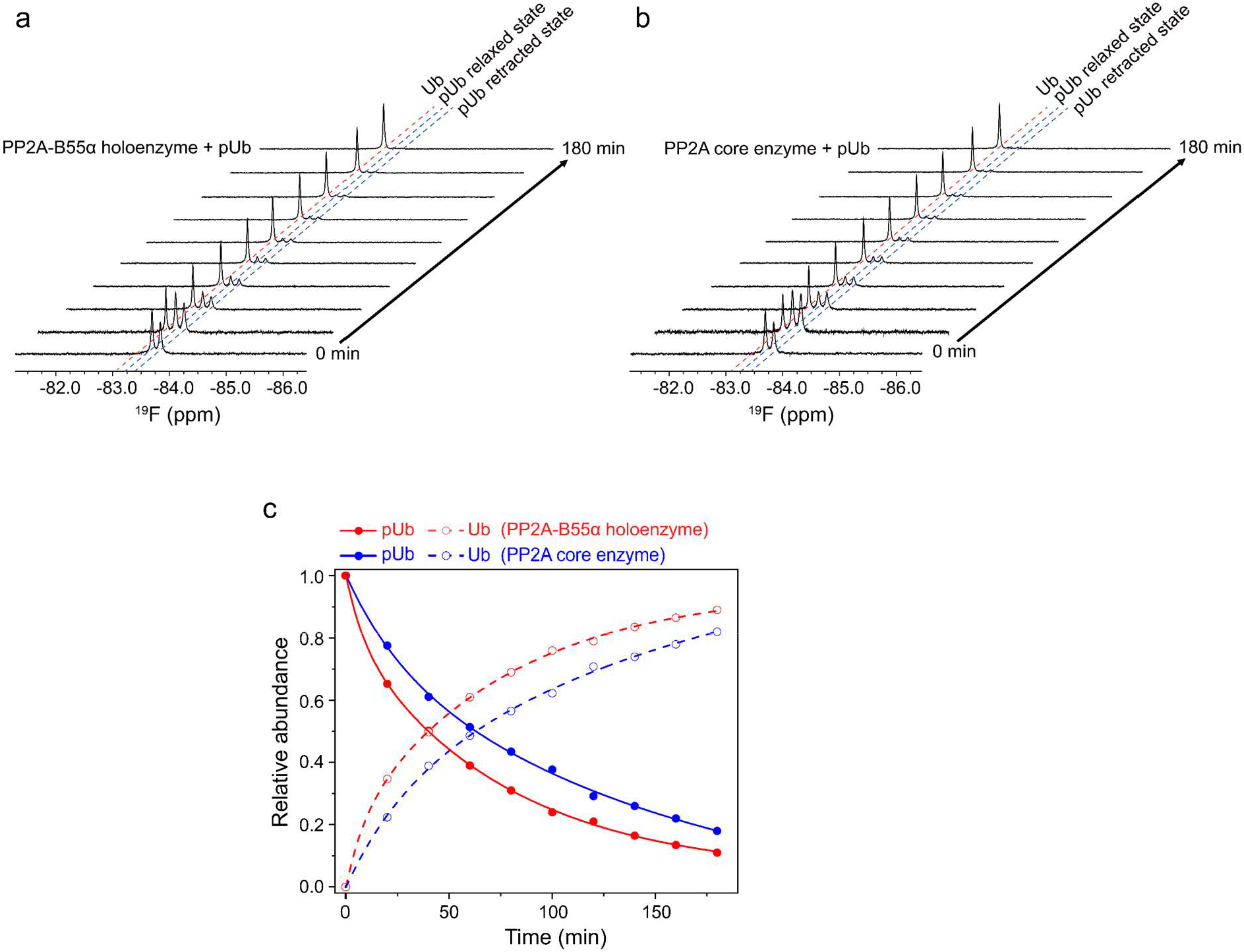
^19^F NMR analysis confirms that B55α accelerates PP2A dephosphorylation of pUb. **a** A stack of ^19^F NMR spectra monitoring the dephosphorylation process of pUb in the presence of PP2A holoenzyme. **b** A stack of ^19^F NMR spectra monitoring the dephosphorylation process of pUb in the presence of PP2A core enzyme. **c** Comparison of the dephosphorylation rates of pUb in the presence of PP2A holoenzyme and PP2A core enzyme. Normalized to the ^19^F peak intensities measured in real-time, referencing to the intensities at time 0; the Ub peak intensity was calculated as the total intensity subtracting remaining pUb peak intensity.

